# Extracellular vesicles as predictors of individual response to exercise training in youth living with obesity

**DOI:** 10.1101/2020.11.20.390872

**Authors:** Taiana M. Pierdoná, Alexandria Martin, Patience O. Obi, Samira Seif, Benjamin Bydak, Ashley Eadie, Keith Brunt, Jonathan M. McGavock, Martin Sénéchal, Ayesha Saleem

**Affiliations:** Diabetes Research Envisioned and Accomplished in Manitoba (DREAM) Theme, Winnipeg, MB, Canada; Faculty of Medicine, Department of Pharmacology, Dalhousie University, Saint John, NB, Canada; Department of Pediatrics and Child Health, Max Rady College of Medicine, University of Manitoba, Winnipeg, MB, Canada; Cardiometabolic Exercise and Lifestyle Laboratory, Faculty of Kinesiology, University of New Brunswick, Fredericton, NB, Canada; Applied Health Sciences, University of Manitoba, Winnipeg, MB, Canada; Children’s Hospital Research Institute of Manitoba (CHRIM), Winnipeg, MB, Canada; Faculty of Kinesiology and Recreation Management, University of Manitoba, Winnipeg, MB, Canada

## Abstract

Exercise is associated with various health benefits, including the prevention and management of obesity and cardiometabolic risk factors. However, a strong heterogeneity in the adaptive response to exercise training exists. The objective of this study was to evaluate if changes in extracellular vesicles (EVs) after acute aerobic exercise (AE) were associated with the responder phenotype following 6-weeks of resistance exercise training. This is a secondary analysis of plasma samples from the EXIT trial (clinical trial #02204670). Eleven sedentary youth with obesity (15.7±0.5 years, BMI ≥ 95th percentile) underwent an acute bout of AE (60% heart rate reserve, 45 min). Blood was collected before exercise [at time (AT) 0 min], during [AT15, 30, 45 min], and 75 min after exercise [AT120]. Afterward, youth participated in 6-week resistance training program, and were categorized into responders (RE) or non-responders (NRE) based on changes in insulin sensitivity as measured by the Matsuda Index. EVs were isolated using size exclusion chromatography (Izon®). The primary outcome variable was EV biophysical profile, which includes size, zeta potential, protein yield and expression of markers associated with EV subtypes. The variables were analyzed in a single-blind fashion. Overall, there was a general increase in EV production in both groups. Average EV size was larger in RE (~147 nm) vs. NRE (~124 nm; p<0.05). Average EV size at AT0 was associated with absolute change in Matsuda index following 6-weeks of resistance training (r=0.44, p=0.08). EV size distribution revealed RE preferentially expressed EVs between 150 – 250 nm in size, whereas NRE expressed EVs between 50 – 100 nm (p<0.05). At baseline, RE-EVs contained ~25% lower Tsg101 protein, ~85% higher MMP2 content, while CD63 levels remained unchanged between the groups. Total protein yield in RE-EVs was higher than NRE at AT15 (p<0.05). Our data suggest that youth with obesity that respond to exercise training produce larger EVs, with lower exosome- and higher microvesicle-specific protein expression. RE-EVs also had higher EV protein yield during AE. The relationship between larger EV subtypes and/or cargo, and the individual response to exercise has yet to be fully elucidated.

## Introduction

Obesity, type 2 diabetes, and hypertension are the most common cardiometabolic diseases in adolescents (1) and are increasingly prevalent worldwide (2). In Canada, nearly 1 in 7 children and youth are overweight or obese (4, 5). With increasing rates of cardiometabolic diseases in youth and adolescents, understanding the impact of prevention strategies is needed, particularly, who responds well to specific intervention types (6, 7). Exercise is fundamental in the management of cardiometabolic risk associated with obesity in youth as it promotes visceral fat loss and cardiorespiratory fitness increases (8, 9). Despite the general effectiveness of exercise on cardiometabolic risk in youth, our group and others have documented notable heterogeneity in the individual response to exercise training. Within clinical trials for example, participants who display positive metabolic improvement post exercise-training are classified as exercise responders, and make up 40-70% of youth in these trials (10). Variable adaptation in the metabolic response to exercise e.g. change in adiposity or hepatic triglycerides content, is linked with the differential increase in cardiorespiratory fitness with training (11). As genetic factors account for ~30-50% of this heterogeneity in response (12), it suggests that other physiological factors likely play a role in this variable response to physical activity.

Chronic exercise training activates a program of transcriptional and non-transcriptional signalling cascades in skeletal muscle that leads to an increase in mitochondrial biogenesis and ultimately metabolic capacity (13). This in turn promotes improved insulin signalling, decrease in weight loss, and enhanced cardiorespiratory fitness (14). Foundational work by Pedersen and colleagues established the role of myokines in potentiating the systemic benefits to regular exercise training (15). Myokines are proteins or peptides released by skeletal muscle into the circulation at rest, and at elevated rates upon exercise (16). Myokines are important in mediating the whole body metabolic response to physical activity (17–20), and have been found packaged within extracellular vesicles (EVs). Little information exists on the interaction between myokines, EVs and their role in regulating the physiological responses to exercise in youth living with obesity.

EVs are small membrane-bound vesicles produced by cells, contain molecular cargo representative of the cell of origin (21), and play a vital role in intracellular communication (22–24). EVs can be present in all biological fluids such as milk, plasma, serum and saliva. EVs are characterized according to size, density, biochemical composition or cell of origin (28). EVs are largely divided into three main subtypes: 1) exosomes (30-150 nm), 2) microvesicles (100-1000 nm) and 3) apoptotic bodies (500-5000 nm). The biogenesis of EV subtypes, their size and biophysical characteristics, and molecular cargo are distinct from one another and likely affect recipient cells differentially upon vesicle uptake. Unless specific markers of EV subcellular origin can be established, according to the Minimal Information for Studies of Extracellular Vesicles (MISEV) 2018 guidelines, EVs should be classified as small EVs (sEVs <200 nm) or medium/large EVs (m/lEVs >200 nm) (29). Recently, studies have shown that exercise promotes the release of EVs from skeletal muscle, platelets, endothelial cells, and leukocytes into the bloodstream (25–27). There is a growing body of evidence that EVs play a role in regulation of the cardiometabolic adaptations to exercise training.

Exercise promotes the release of EVs from skeletal muscle, platelets, endothelial cells, and leukocytes into the bloodstream (24–27). Given this, we previously hypothesized that EVs may play a functionally important role in determining the systemic metabolic adaptations to training (24). However, we do not know if the variance in the cardiometabolic response to exercise is associated with changes in EV subtypes and biophysical characteristics. To address this gap, we performed a secondary analysis of plasma samples from the EXIT study (Clinical Trials Number: 02204670, (30)]. The EXIT study investigated the influence of the exercise-induced myokine, irisin, on the metabolic response to exercise training in obese youth (30). Our main objective was to determine if EVs released during an acute bout of aerobic exercise (AE) prior to 6-weeks of resistance training were associated with the exercise responder phenotype. Within this secondary analysis of data the EXIT trial, we hypothesized that obese youth that are exercise responders will be characterize by biophysically different EVs (by size, yield, and/or stability) as well as by molecular cargo.

## Materials and methods

### Participants and plasma

Participants underwent a baseline screening, which included medical history and an oral glucose tolerance test and a baseline assessment, which included assessment of body composition, fitness, physical activity level, and muscle strength. Following the first visit, youth completed an acute bout of aerobic exercise (AE), during which blood draw were taken. Following those two acute exercise sessions, all participants performed a 6-week resistance exercise intervention to determine the metabolic response to chronic resistance exercise training. Plasma samples that were available at various time points from the EXIT trials [Clinical Trials Number: 02204670, (30)] were available from all youth participants (n=11). We isolated EVs from the plasma samples that were collected before, during, and after an acute bout of AE by cycling at 60% heart rate reserve for 45 min. Blood was collected through a venous catheter from an antecubital vein by a registered nurse, before AE [acute training time (AT) −15, 0 min], during AE [AT15, 30, 45 min], and after AE + recovery [15 min (AT60), 75 min (AT120)]. Blood was centrifuged and plasma collected and stored at −80°C for further analysis. Afterward, youth participated in 6-week resistance exercise training program that consisted of 8 different exercises performed (3 sets of 12–15 repetitions at 60–65% of the estimated 1-max repetition). The study was approved by the University of Manitoba Biomedical Research Ethics Board and performed according to the Declaration of Helsinki.

### EV isolation

EVs were isolated using size exclusion chromatography (SEC) (qEV single columns, Izon Science Ltd, New Zealand) according to manufacturer’s instructions. We extracted EVs from plasma samples collected at baseline (AT0), during AE (AT 15, 30 and 45) and during recovery (AT120) as shown in Figure 1A. Briefly, plasma samples were passed through a 0.22 μm filter (Merck Millipore, MA, USA) to eliminate cells and cellular debris. Next, 150 μL of each sample was loaded into qEV columns and 12 x 200 μL fractions (F) were collected with filtered phosphate-buffered saline (PBS) as the elution buffer. Each fraction (F1–12) was analyzed for EV size, protein yield and presence of EV specific markers as well as negative controls by Western blotting. F7–10 were considered EV-rich, plasma protein- and lipoprotein-poor, and were pooled to measure EV size, zeta potential, total protein yield and presence of EV subtype markers by Western blotting.

**Figure 1.**
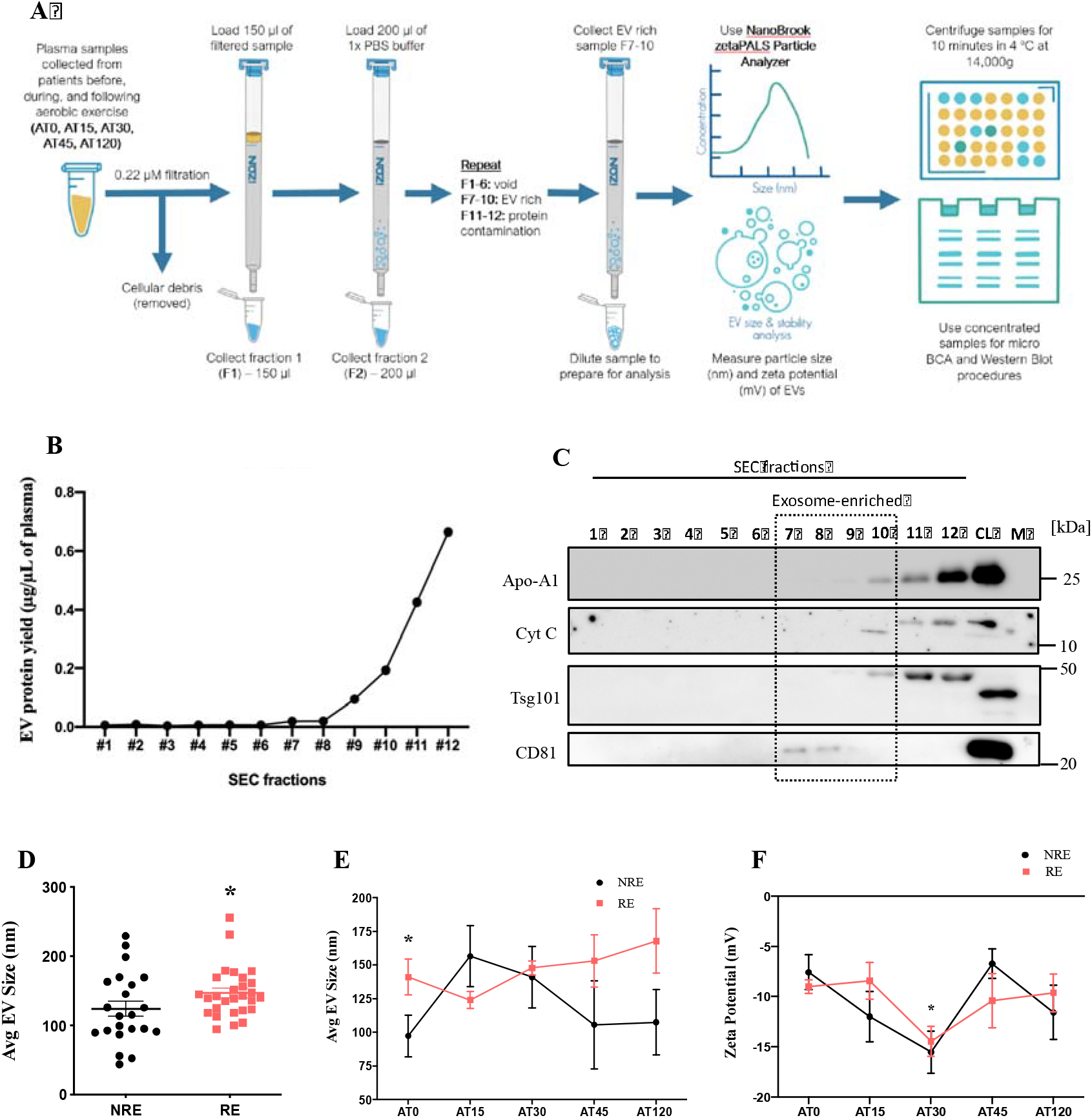
Study design, SEC validation and biophysical characterization of EVs from EXIT trial participants. **(A)** Schematic depicts the study design. Plasma was collected from participants in the EXIT trial study, before (AT0), during (AT 15, 30 and 45) and after recovery (AT120) of AE training. EVs from non-responders (NRE-EV) and exercise responders (RE-EV) were isolated by SEC and analyzed for size and zeta potential using DLS (NanoBrook ZetaPALS). **(B)** The protein yield was measured using Pierce™ MicroBCA Protein Assay kit and showed an exponentially increasing protein concentration from fraction 7 (F7) onwards. **(C)** Western blotting was performed (12% SDS-PAGE) and Coomassie blue staining used as a loading control. Proteins enriched in exosomes (TSG101 and CD81), cytochrome c (Cyt. C, negative control i.e. non-exosomal protein), and Apolipoprotein A1 (ApoA1) as lipoprotein marker were quantified. Fractions 7 to 10 were considered exosome-rich while lipoprotein-poor. **(D)** Average EV size was larger in RE-EV group compared with NRE-EV overall, **(E)** and especially at AT0 (*p<0.05). **(F)** Zeta potential was unchanged between the two groups and decreased in both at AT30 (*p<0.05). Data were analyzed using multiple unpaired Student’s t-test, or with *p<0.05 considered as significant and expressed as mean ± standard error (N=5-6).

### EV characterization

EVs were stored at 4 °C for up to 24 hours before measuring size and zeta potential using a NanoBrook ZetaPALS (Brookhaven Instruments). Prior to characterization, 20 μl of the fraction was diluted 1:75 in PBS and dilutions kept on ice until analysis. Size distribution was measured through analysis of multimodal size, and size bins were used to represent total size intensity within a given size range. For zeta potential, each sample was loaded into a disposable cuvette, and an electrode was inserted. Each sample underwent 5 runs that were averaged (irrespective of negative/positive charge) to calculate zeta potential using the Smoluchowski formula (31).

### Protein concentration and quantification

To concentrate the protein content in the samples, F7–10 were centrifuged at 14,000x g, at 4°C for 60 min using an Amicon Ultra-4 3 kDa centrifugal filter device (Merck Millipore). The 50 μL retentate was the final volume of concentrated EVs. Proteins were extracted using 1:1 Pierce RIPA solution with protease inhibitor tablet (Roche). Samples were vortexed for 5 seconds, followed by two freeze/thaw cycles with liquid nitrogen, sonicated 3×3 seconds on ice and centrifuged at 15,000 rpm (21,130x g) for 15 minutes at 4°C. Proteins were quantified using the Pierce™ Micro BCA Protein Assay kit (Thermo Scientific) following the manufacturer’s instructions. Briefly, the working reagent was prepared by mixing 25 parts of Reagent A, 24 parts of Reagent B and 1 part of Reagent C. The standards were prepared by serial dilution of 2 mg/ml bovine serum albumin (BSA) ampule into clean vials using ultrapure water. 150 μL of each standard or sample was added to a 96-well microplate in duplicates, followed by the addition of 150 μL of working reagent to each well. The samples were incubated 37 °C for 2 hours and absorbance was measured at 562 nm using a microplate spectrophotometer (BioTek Epoch).

### Western blotting

Western blotting was used to determine the expression of subtype specific protein expression in order to ascertain the subcellular origin of EVs. Samples were denatured by the addition of β-mercaptoethanol and 4x laemmli buffer, followed by incubation at 95 °C for 5 min. Then, 7 μg of total protein was loaded on 12% SDS-PAGE gels and electrophoresed at 90 mV for 30 min and then 120 mV for 2 hours. Proteins were transferred to polyvinylidene difluoride (PVDF) membrane using Trans-Blot® Turbo™ (Bio-Rad). Next, the membrane was incubated with blocking buffer [5% skim milk in Tris-buffered saline Tween-20 solution (TBST)] for 1 hour at room temperature. The PVDF membrane was incubated with the primary antibodies in 1% skim milk overnight, at 4 °C. The following primary antibodies were used: rabbit polyclonal anti-TSG101 (T5701, Sigma-Aldrich Co, 1:200), rabbit polyclonal anti-CD63 (SAB4301607, Sigma-Aldrich Co, 1:300), mouse monoclonal anti-CD81 (sc-166029, Santa Cruz Biotechnology, 1:200), rabbit polyclonal anti-Cytochrome C (Cyt C) (AHP2302, Bio-Rad Laboratories, 1:500), mouse monoclonal anti-MMP2 (sc-13595, Santa Cruz Biotechnology, 1:200), mouse monoclonal anti-Apolipoprotein A1 (Apo-A1) (0650-0050, Bio-Rad Laboratories, 1:200). Membranes were washed 3 times with TBST and incubated with anti-mouse IgG secondary antibody (A16017, Thermofisher, 1:10,000) or anti-rabbit IgG secondary antibody (A16035, Thermofisher, 1:1,000) for 1 hour at room temperature. The Clarity™ and Clarity Max™ ECL substrate (Bio-Rad) was applied to visualize bands via enhanced chemiluminescence and images captured using ChemiDoc^TM^ MP Imaging System (Bio-Rad). Band intensity was quantified using Image lab software (Bio-Rad) and corrected for loading using Coomassie blue dye staining.

### Resistance exercise training

The resistance training program has been describe elsewhere. Briefly, the 6-week resistance training program, consisted of three weekly sessions on non-consecutive days conducted at the local YMCA in Winnipeg. Participants warmed up on a cycle ergometer, treadmill or an elliptical for about 5-10 minutes before each session. Resistance training consisted of 8 different exercises including seated chest press, leg extension, narrow grip latissimus pull down, seated leg curl, arm curl, shoulder elevation, triceps extension, and the plank. Each exercise was performed for 3 sets of 12-15 repetitions at 60-65% of the estimated one-repetition-maximum (1-RM). Resting periods of 60 seconds separated each set.

### Definition of responder or non-responder to exercise training

Participant performed an oral glucose tolerance test (OGTT) at baseline and post intervention. Plasma glucose and insulin were used to determine the Matsuda index, calculated from the glucose and insulin response to a frequently-sampled oral glucose tolerance test. Blood samples were taken at baseline, 30, 60, and 120 minutes following the ingestion of a 75-gram glucose drink. Matsuda index according to the following formula: 10 000/ √ (Fasting Plasma Glucose x Fasting Plasma Insulin) x (G x I). Following the intervention, participants were categorized into responders to exercise (RE) if change in insulin sensitivity was above the 50 percentile, or non-responders to exercise (NRE) if change in insulin sensitivity was below the 50 percentile.

### Statistical analysis

Average EV size, zeta potential, size distribution, protein yield, and expression of protein markers at AT0, data were analyzed using multiple unpaired Student’s t-test. A one-way ANOVA with Tukey post-hoc was used to analyze protein expression over time, with values expressed as fold change over AT0. Average EV size pre-training and absolute change in Matsuda Index post-training were analyzed using Pearson’s Correlation Coefficient. All data were analyzed using PRISM software, version 8.4.2 (GraphPad, San Diego, CA) with 95% confidence intervals. Significance was set at *p<0.05 and data were expressed as mean ± standard error (N=5-6).

## Results

### Biophysical characterization of isolated EVs from responders *vs*. non-responders to exercise

Pooled EV-rich fractions (F7–10) from RE or NRE participants (RE-EV and NRE-EV) isolated via SEC were characterized. EV size (nm) and zeta potential (mV) was measured by dynamic light scattering (DLS) using a NanoBrook ZetaPALS. Samples were concentrated, total protein yield determined and markers of subcellular origin (exosomes/microvesicles) were measured by immunoblotting (**Figure 1A**). Based on its efficiency, reproducibility, dependability, and accuracy, SEC-based approach is an established technique for the isolation of pure, homogenous and intact EVs, and in particular sEVs and exosomes (28). To validate SEC-based EV isolation in our study, as per MISEV guidelines (29) we analyzed each fraction (F1–12) for protein yield and expression of exosome-specific (TSG101 and CD81) or non-exosomal markers or negative controls (Cyt C and Apo-A1). Protein concentration increased nearly exponentially starting in F7 through to F12 (**Figure 1B**) as shown before (32, 33). Exosomal markers TSG101 and CD81 were enriched in F7–10. While, F10 was not completely free of contaminating lipoproteins (ApoA1) or non-EV plasma proteins (Cyt C), the expression of these proteins was lower when compared to F11–12 and cell lysate (CL) (**Figure 1C**). Furthermore, EV size analysis on F10 revealed it contained particles sized 56.3 ± 17.8 nm. This established that F7–F10 are sufficiently enriched with TSG101+/CD81+ sEVs that were subsequently pooled for all further analysis.

Average EV size (all time points pooled) was larger in the RE group (146.9 ± 6.8 nm) compared to the NRE group (124.1 ± 11.0 nm) (*p <0.05, **Figure 1D**). Average EV size over the AE bout (AT15, 30, 45) and recovery (AT120) showed larger EVs in the RE group at nearly all time points except at AT15, where NRE group presented larger particles (156.5 ± 22.7 nm *vs*. 124.0 ± 6.4 nm). Moreover, the groups were significantly different at AT0, demonstrating that the subjects were releasing EVs of different sizes basally even before initiating acute exercise (*p <0.05) (**Figure 1E**). Zeta potential indicative of EV stability in suspension (34), remained comparable between the two groups (NRE: −10.7 ± 1.6 mV and RE: −10.4 ± 1.0 mV). Interestingly, zeta potential was lowest (i.e. EVs were the most stable) at AT30 for both groups (NRE: −15.5 ± 2.1 mV and RE: −14.4 ± 1.5 mV) when compared to the other time points (*p <0.05) (**Figure 1F**).

Overall size distribution across all-time points re-affirmed that RE had a higher yield of m/lEVs, between 150 and 250 nm compared with the NRE group. In contrast, the NRE group had elevated levels of sEVs sized 50 to 100 nm (*p <0.05, **Figure 2A**). Analyses of EV size distribution at baseline (AT0), during exercise (AT15 to AT45) and recovery (AT120) showed a similar pattern where RE group expressed increased m/lEVs, whereas the NRE group had mostly sEVs (*p <0.05, **Figure 2B**). Interestingly, NRE-EV size distribution matched the pattern of the RE-EV counterparts and both showed an increased expression of m/lEVs at AT30 (**Figure 2B**). Size distribution analysis demonstrated a general increase in EV production with exercise and recovery in both groups, when results were expressed for each group individually (**Figure 2C**). The pattern of preferentially higher expression of m/lEVs in responders to exercise, and sEVs in non-responders remained true irrespective of exercise or recovery (**Figure 2C**).

**Figure 2.**
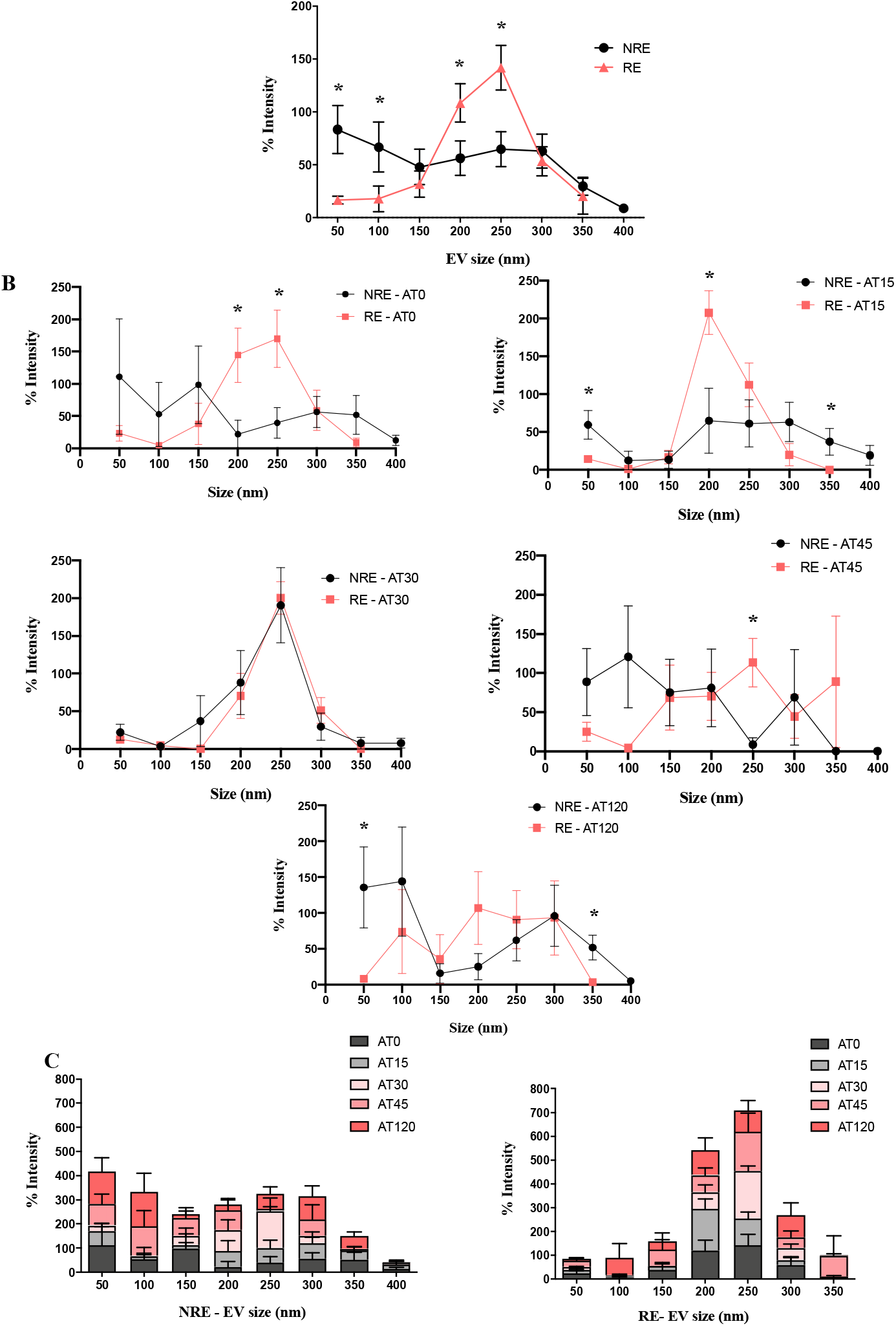
EV size distribution profile in responders *vs*. non-responders to exercise. **(A)** EV size distribution analysis combined for all time points illustrated a higher yield of medium/large EVs (m/lEVs) in the RE group between 150 nm to 250 nm. NRE group expressed increased content of small EVs (sEVs) between 50 nm to 100 nm size range (*p<0.05). **(B)** Analyses of EV size distribution by individual time points showed RE group with an increase in m/lEVs at baseline (AT0) and during AE (AT 15, 30 and 45), while NRE showed enhanced expression of sEVs at all time points except AT30 where it matched the RE group (*p<0.05). **(C)** Size distribution demonstrated a general increase in EV production with time in both groups. Data were analyzed using multiple unpaired Student’s t-test with *p<0.05 considered as significant and expressed as mean ± standard error (N=5-6).

### Western blot analysis of proteins enriched in EV subtypes

Total protein yield of EVs isolated from both groups remained unchanged at baseline (AT0). The protein yield for the RE group remained consistent across the study, but the NRE group showed a significant decrease in protein content during the first 30 min of AE (NRE: 0.08 ± 0.03 μg/μL *vs*. RE: 0.25 ± 0.03 μg/μL at AT15) (*p <0.05, **Figure 3A**). This difference diminished at AT30 and remained as in the initial state at AT45 and AT120 (**Figure 3A**). Western blot analysis was performed to assess expression of exosome and microvesicle-specific proteins in line with MISEV guidelines (29). We examined the expression of the transmembrane tetraspanin cluster of differentiation 63 (CD63) and a multivesicular body marker tumor susceptibility gene 101 (TSG101) as representative proteins enriched in exosomes. To investigate the presence of microvesicles, we measured expression of matrix metalloproteinases 2 (MMP-2). RE-EVs expressed ~25% lower TSG101 and ~85% higher MMP2 content, while CD63 levels remained unchanged between the groups at AT0 (**Figure 3B**). Next we assessed the expression of CD63, TSG101 and MMP-2 over time (AT15, 30, 45 and 120) in both groups. The expression of exosomal and microvesicle-related proteins with AE in both groups were statistically insignificant, likely due to the low sample size (**Figure 4**).

**Figure 3.**
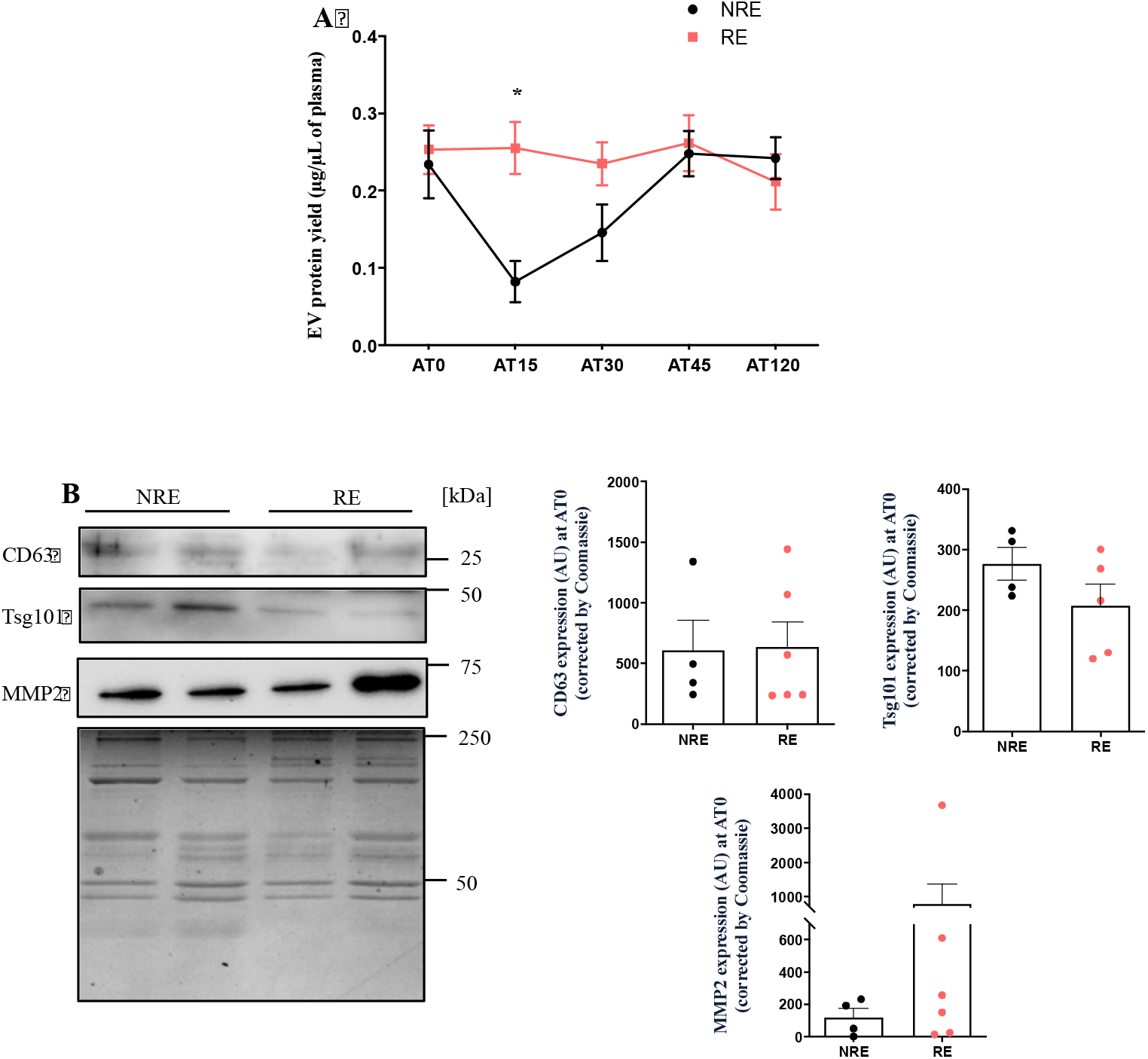
Protein yield and expression of markers of EV subtypes AT0. **(A)** NRE-EV group showed a significant decrease in protein yield compared to RE-EV at AT15 (*p<0.05). **(B)** Equal amounts of NRE-EV or RE-EV protein (7 μg/mL) were subjected to SDS-PAGE (12%) and expression of proteins enriched in exosomes: TSG101 (46kDa) and CD63 (28kDa), and microvesicles: MMP2 (63kDa) was quantified. RE-EVs expressed ~25% lower Tsg101 protein, ~85% higher MMP2 content, while CD63 levels remained unchanged between the groups at AT0. Coomassie staining was used as a loading control. Data were analyzed using unpaired Student’s t-test with *p<0.05 considered as significant and expressed as mean ± standard error (N=5-6).

**Figure 4.**
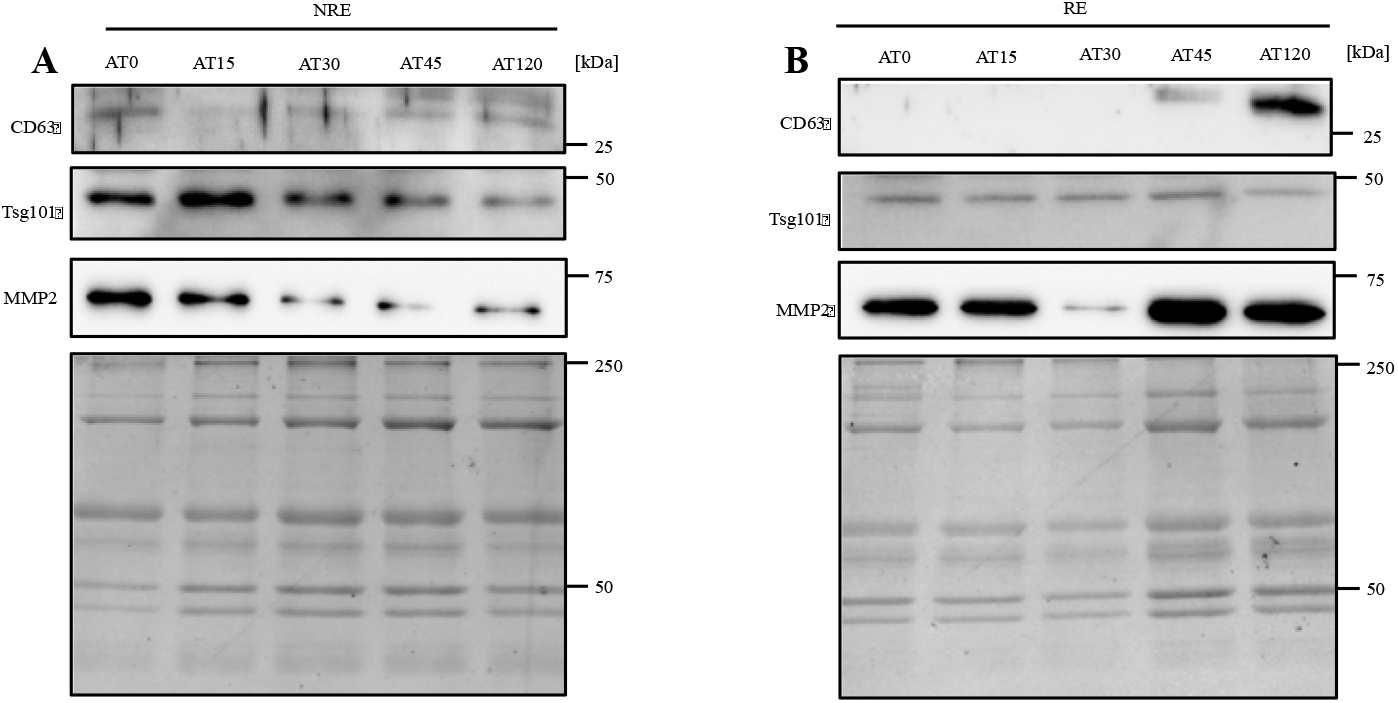
Expression of exosome and microvesicle markers at baseline, during and after exercise. **(A)** Representative blots for NRE-EV and **(B)** RE-EV showing expression of exosome (TSG101, CD63) and microvesicle (MMP2) proteins at all time points of the AE. Coomassie staining was used as a loading control. No significant changes were observed for any protein. Data were analyzed using one-way ANOVA (N=3-4).

### EV size and its correlation with insulin sensitivity

To investigate if there is any correlation between EV size and insulin sensitivity in the participants, we applied the Pearson’s correlation coefficient to our data. Our results suggested that there is a moderate positive correlation between average EV size and absolute change in Matsuda Index post-training (r=0.4374, p=0.08) (**Figure 5**).

**Figure 5.**
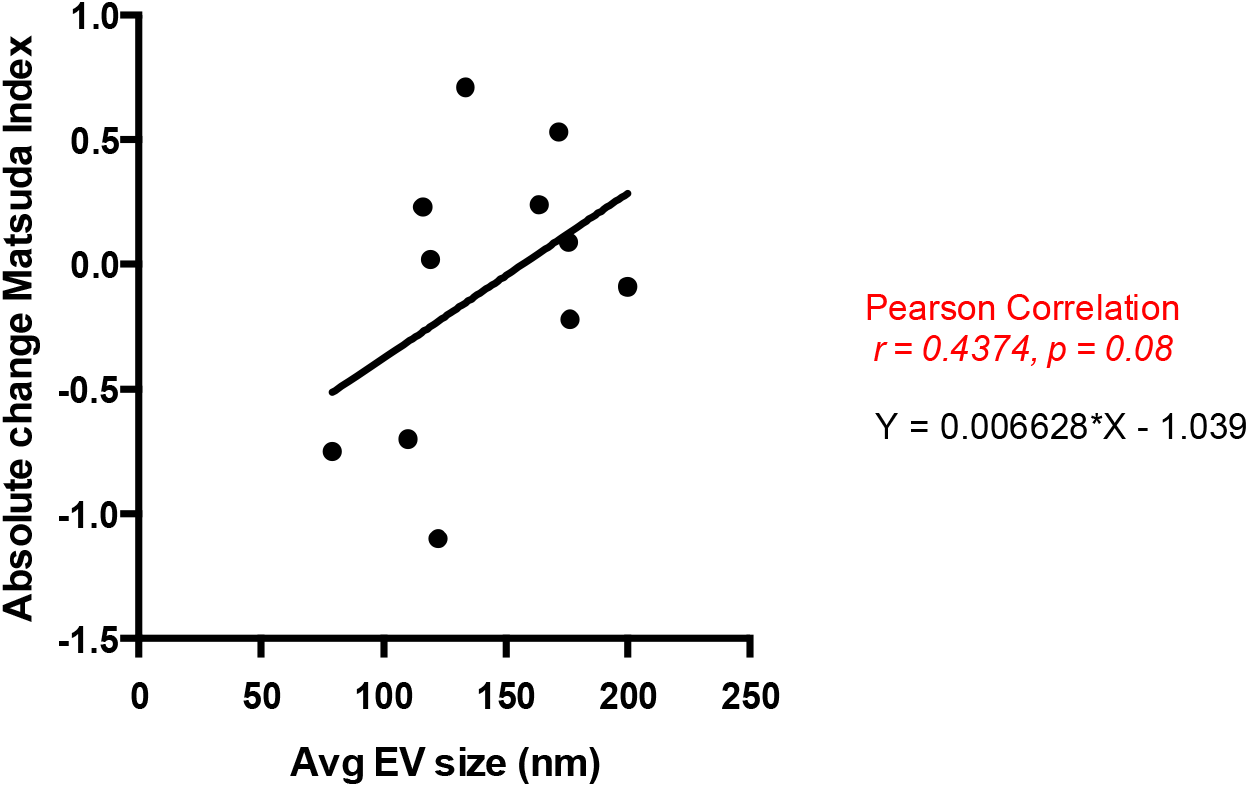
Association of EV size with absolute change in Matsuda Index. Pearson’s Correlation analysis showed a moderate positive association between average EV size at AT0 and absolute change in Matsuda Index (indicative of insulin sensitivity) post resistance exercise training (r=0.4374, p=0.08).

## Discussion

In the current study we sought to elucidate whether changes in EV biophysical characteristics (size, yield) were associated with an individual’s capacity to respond to exercise training in a single-blind fashion. The main findings of our study are: (1) acute exercise leads to an increase in EV yield in participants; (2) average EV size was larger in responders (RE; 150 – 250 nm), *vs*. non-responders (NRE, 50 – 100 nm); and (3) the size of EVs was aligned with expression of markers enriched in EV subtypes, microvesicles *vs*. exosomes. The RE group produced EVs with lower expression of exosomal proteins and higher expression of microvesicles markers. Lastly, we also observed a mild positive correlation between overall average EV size and the responder phenotype. These results are important as they shed light into the potential mechanism of exercise response in obese youth at risk of Type 2 diabetes.

Previous studies have shown that circulating EVs are elevated in obese mice and human (41, 42), however, the potential mechanisms that lead to higher EV release and the biological effects of the EV-subtypes in obesity are not yet known (43). Our data showed different populations of EVs in the RE and NRE groups where the size distribution was significantly larger in RE (150 – 250 nm) when compared to the NRE (50 - 100 nm) group. Durcin et al. (2017) showed that adipocytes release different populations of EVs, including lEVs and sEVs (38), and express distinct proteins enriched in microvesicles and exosomes, that predict their biological function. Others have demonstrated that white adipose tissue (WAT)-derived EVs hold immunomodulatory properties and can modulate insulin signalling in muscle and liver cells (44). Thus, both EV size and cell of origin can affect downstream functional effect of EVs on recipient cells. We propose that, the functional responses of EVs from RE *vs*. NRE may be an underlining mechanism of differential improvements in insulin sensitivity in the participants.

Our study demonstrated a general increase in EV production over time in both groups after AE. This is in agreement with previous work that has demonstrated an increase of EVs in healthy (25, 45) as well as middle-aged, overweight individuals (23) after AE. Whitman and colleagues (2018) showed that one hour of cycling induced a significant increase in systemic EVs (45), while Frühbeis and collaborators (2015) reported that the systemic EV concentration was higher immediately after a single exhaustive resistance exercise (25). Another study showed that acute AE was associated with an increase in EV concentration and the presence of twelve different miRNA in their cargo (46). In contrast, in obese adults, Rigamonti et al. (2019) reported reduced EVs concentration immediately at the end of AE and after 3h and 24 h (47). No studies have been conducted on EV release with AE in youth with obesity to our knowledge. Other than the obvious difference in participant age, the level of circulating EVs is dynamic and can be modulated by a number of factors including body weight (42), age, immune status, hormone levels and metabolic state (48) that can explain the discrepancy in our result *vs*. previous work. Lastly, the AE time course in our study is much shorter compared to Rigamonti et al. who used measured EVs immediately after, 3h and 24h later. Further studies are needed to understand the temporal pattern of EV release with exercise in healthy adults and youth, as well as those with metabolism-related conditions such as obesity, type 2 diabetes and the metabolic syndrome.

EVs released after AE may originate from cell types of the circulatory system such as platelets, endothelial cells and leukocytes, as well as skeletal muscle (45, 48). A recent study (49) showed that obese individuals with very poor fitness presented elevated EVs of platelet and endothelial origin versus people with poor fitness, independent of age and body fat. The authors suggested that subtle differences in fitness may reduce type 2 diabetes and cardiovascular disease risk through an EV-related mechanism. Due to limited samples, we were unable to ascertain the origin of the EVs isolated in our study, but this research is necessary to understand the origin and possible downstream target of EVs in youth living with obesity.

There is growing recognition of EVs as biomarkers of cardiometabolic diseases (35–37), and we were one of the first group to hypothesize that endurance exercise-derived exosomes will rescue metabolic diseases (24, 35–37). However, these findings need to be interpreted carefully due to the technical limitations associated with the different EV isolation and characterization techniques, and the heterogeneity of EV subtypes and their biological significance (38, 39). Previous studies showed that SEC minimally alters the characteristics of isolated EVs and in particular, exosomes. SEC is considered to be one of the best methods for separating exosomes from protein contaminants and co-precipitates (28, 40). As per MISEV guidelines, we first measured the purity of isolated EV fractions. We showed an increase in the expression of proteins enriched in exosomes, TSG101 and CD81 in F7–10 as reported previously (32, 33). While F10 was not completely free of contamination by lipoproteins (ApoA1) and non-exosomal proteins (Cyt C), it also showed the presence of sEVs (~56.3 nm, data not shown). Our results are compatible with other studies that reported similar findings when using SEC with qEV columns (32, 33).

RE showed significantly higher concentrations of total protein yield in isolated EVs than the NRE group. This was manifest due to the sharp decrease in the protein yield in the NRE group after starting the exercise, particularly evident at AT15. This is in line with Oliveira et al. (2018) who found that AE significantly increased serum EV protein yield in rats (46). Durcin et al. (2017) also reported that despite the reduced secretion of large EVs by 3T3-L1 adipocytes, a higher diversity of proteins was identified in the larger EV fractions (38). We showed in our study that EVs released at AT15 and AT30 had lower zeta potential values (i.e. more stable particles) in both groups and therefore the protein content may be better conserved in this condition. Furthermore, we infer that during the AE, the RE group was able to release more proteins packaged in lEVs when compared to the NRE group and these could encapsulate myokines. However, more studies are needed to identify the EV cargo proteins and their main role in the pro-metabolic exercise response to AE.

While both RE and NRE groups expressed proteins enriched in exosomes (TSG101 and CD63) as well as microvesicles (MMP-2), we measured 25% higher TSG101 expression in NRE, and ~85% higher MMP2 content in RE group. Our data corroborate previous findings (28) of enriched exosomal markers Alix, TSG101 and tetraspanins (CD9, CD63 and CD81) in sEVs. Interestingly, previous studies report that lEVs carry proteins of different origins and are specifically enriched in metabolic enzymes (mainly of mitochondrial origin), suggesting that lEVs may modulate metabolic pathways, such as fatty acid oxidation (38). We suspect that the RE group that produced lEVs adapted to the exercise because of the metabolic EV cargo content in lEVs. Targeted experiments to specifically isolate lEVs only and deduce the effect on metabolic adaptations are warranted based on these outcomes. Finally, while the EXIT trial study (30) showed a correlation between absolute change in irisin and insulin sensitivity following 6-weeks of resistance training, we found that overall average EV size is moderately associated with improved insulin sensitivity.

Our study provides information that youth living with obesity who responded to exercise produced EVs that are larger in size, with a higher protein yield, lower expression of exosomal proteins and higher expression of microvesicles, *apriori* to exercise training. We do not know the impact of mostly sEVs, which are usually associated with exosomes, in the NRE group. The relationship between the EV cargo and an individual’s ability to respond to exercise has yet to be fully elucidated. We also documented a unique temporal pattern of EV release during exercise and immediately after recovery in both groups. Our results highlight the need to distinguish EV subtypes to delineate their respective functional properties and subsequent effect on response to exercise training.

## Author Contributions

T.M.P and A.M. performed most of the experiments in the current study and wrote the manuscript. P.O.O, S.S, B.B. provided technical support and assistance with experiments. A.E., K.B., J.M.M., M.S. conducted the original EXIT trial clinical study. All authors were involved in manuscript revisions. A.S. designed the project, and helped write the manuscript. A.S. is the corresponding author and directly supervised the project. All authors have read, edited and agreed to the published version of the manuscript.

## Acknowledgments

We would like to thank Dr. Hagar Labouta for use of the Nanobrook zetaPALS.

